# Fatty acids from adipocyte lipolysis stimulate insulin secretion

**DOI:** 10.64898/2026.05.13.724851

**Authors:** Camille Fournes-Fraresso, Emilie Courty, Ebru Temiz, Marie Marques, Stéphanie Cassant-Sourdy, Laura Reininger, Anouk Pellerin, Laure Rolland, Ayse S. Dereli, Etienne Mouisel, Vincent Poitout, Matthieu Raoux, Patrick Gilon, Jean-Sébastien Annicotte, Dominique Langin, Pierre-Damien Denechaud

## Abstract

White adipose tissue and pancreatic islets play central roles in the regulation of metabolic homeostasis. Although ectopic lipid accumulation is established as a driver of impaired insulin secretion, the acute contribution of adipocyte lipolysis to islet function remains poorly documented. Here, we investigated a mouse model with inducible adipocyte-specific deletion of both adipose triglyceride lipase (ATGL) and hormone-sensitive lipase (HSL), which leads to defective adipocyte lipolysis. Despite preserved ex vivo islet function, these mice displayed a marked reduction in insulin secretion in response to stimulation of adipocyte β_3_-adrenoceptors, as well as following glucose and arginine challenges. Mechanistically, we identified non-esterified fatty acids as critical mediators of lipolysis-driven insulin secretion, engaging pancreatic signaling of the free fatty acid receptors FFAR4 (a.k.a. GPR120) and FFAR1 (a.k.a. GPR40). The regulation of insulin secretion by adipocyte lipolysis was preserved in high-fat diet-induced obesity. These findings identify an underappreciated adipose–islet crosstalk that couples adipocyte lipolysis to insulin secretion and links lipid and glucose metabolism.

## INTRODUCTION

In mammals, glucose homeostasis is regulated by a complex network of metabolic processes involving a variety of factors, in which pancreatic hormones are critical^1,2^. Insulin secretion from pancreatic islet β cells is essential for adapting blood glucose levels to environmental changes. In contrast to glucagon produced from α cells, insulin suppresses hepatic glucose production and stimulates tissue glucose uptake. Defective insulin secretion can lead to the development of diabetes mellitus and other metabolic complications^1,2^. The main stimulus for insulin secretion is the increase in circulating glucose in the postprandial state. The effect of glucose on insulin secretion is crucially modulated by a large number of inputs, including nutrients such as amino acids^3^ and free fatty acids^4,5^.

Fatty acids may act directly on β cells to induce insulin secretion via the fatty acid receptor FFAR1 a.k.a. GPR40, or by mobilizing islet intracellular lipids^6–8^. Intracellular lipolysis and glycerolipid cycle in β cells sustain insulin secretion^9^. In addition, evidence suggests that fatty acids enhance both insulin and glucagon secretion by relieving somatostatin inhibitory signaling^6,10^. Indeed, fatty acid activation of pancreatic δ-cell FFAR4 (a.k.a. GPR120) inhibits somatostatin secretion and paracrine action on somatostatin receptors localized on the surface of α and β cells^10–12^.

Multiple sources of fatty acids can modulate pancreatic β-cell function^2,6^. Fatty acids may derive from the diet through the hydrolysis of lipoprotein triacylglycerols (TG) or from intra-islet TG stores^6,8^. In addition, they can come from the mobilization of the body’s largest reservoir of TG through adipose tissue lipolysis which constitutes the major source of circulating non-esterified fatty acids (NEFA)^13^. The latter suggests an inter-organ interplay between adipocytes and pancreatic islets that remains largely under-explored^14–16^, due at least in part to a lack of appropriate genetic tools targeting adipose tissue lipolytic enzymes. Identifying such modulators of insulin secretion could improve our understanding of the regulation of glucose homeostasis and identify new targets to improve insulin secretion.

Adipose lipolysis is a fine-tuned catabolic process regulated by lipolytic (e.g. β-adrenergic) and antilipolytic (e.g. insulin) signals. It increases during fasting to meet the body’s energy demands. Catecholamines binding to adipocyte β-adrenergic receptor (β-AR) promotes cAMP-dependent protein kinase activation which results in phosphorylation of downstream players acting at the surface of adipocyte lipid droplets^13^. Adipose triglyceride lipase (ATGL encoded by *Pnpla2*) and hormone-sensitive lipase (HSL encoded by *Lipe*) are the rate-limiting enzymes responsible for the sequential breakdown of TGs^13^. In rodents, the β_3_-AR, which is predominantly expressed in adipocytes, is the primary mediator of lipolysis^13^. Administration of a β_3_-AR agonist increases insulin secretion^17,18^. In dogs, infusion of long-chain fatty acids into the pancreatic artery stimulates insulin release without significantly increasing peripheral plasma NEFA^4,5^. In humans, treatment with nicotinic acid lowers both plasma NEFA and insulin levels in healthy individuals and type 2 diabetic patients^19^. Despite these investigations, the direct contribution of adipose tissue lipolysis to islet function has not been determined.

We hypothesized that white adipose tissue could directly regulate insulin secretion from pancreatic β cells. To directly address this hypothesis, we generated mice with inducible adipocyte-specific deletion of ATGL and HSL and studied insulin secretion under different drug, substrate and nutritional challenges. To decipher the mechanism of action, we investigated somatostatin- and FFAR4-deficient mice as well as the effect of a FFAR1 agonist. We determined whether NEFA derived from adipocyte lipolysis enhance insulin secretion in lean and obese mice.

## RESULTS

### Impairment of adipocyte lipolysis affect β_3_-AR-, glucose- and arginine-dependent insulin secretion

To investigate the role of adipocyte lipolysis on insulin secretion from pancreatic β cells, we generated a tamoxifen-inducible mouse model deficient for both ATGL and HSL in adipocytes named DaKO (for Double adipo-KnockOut). Following tamoxifen gavage, DaKO mice showed a robust decrease in adipose tissue ATGL and HSL levels compare to sex tamoxifen-treated control littermate (Figure 1A and 1B).

**Figure 1:**
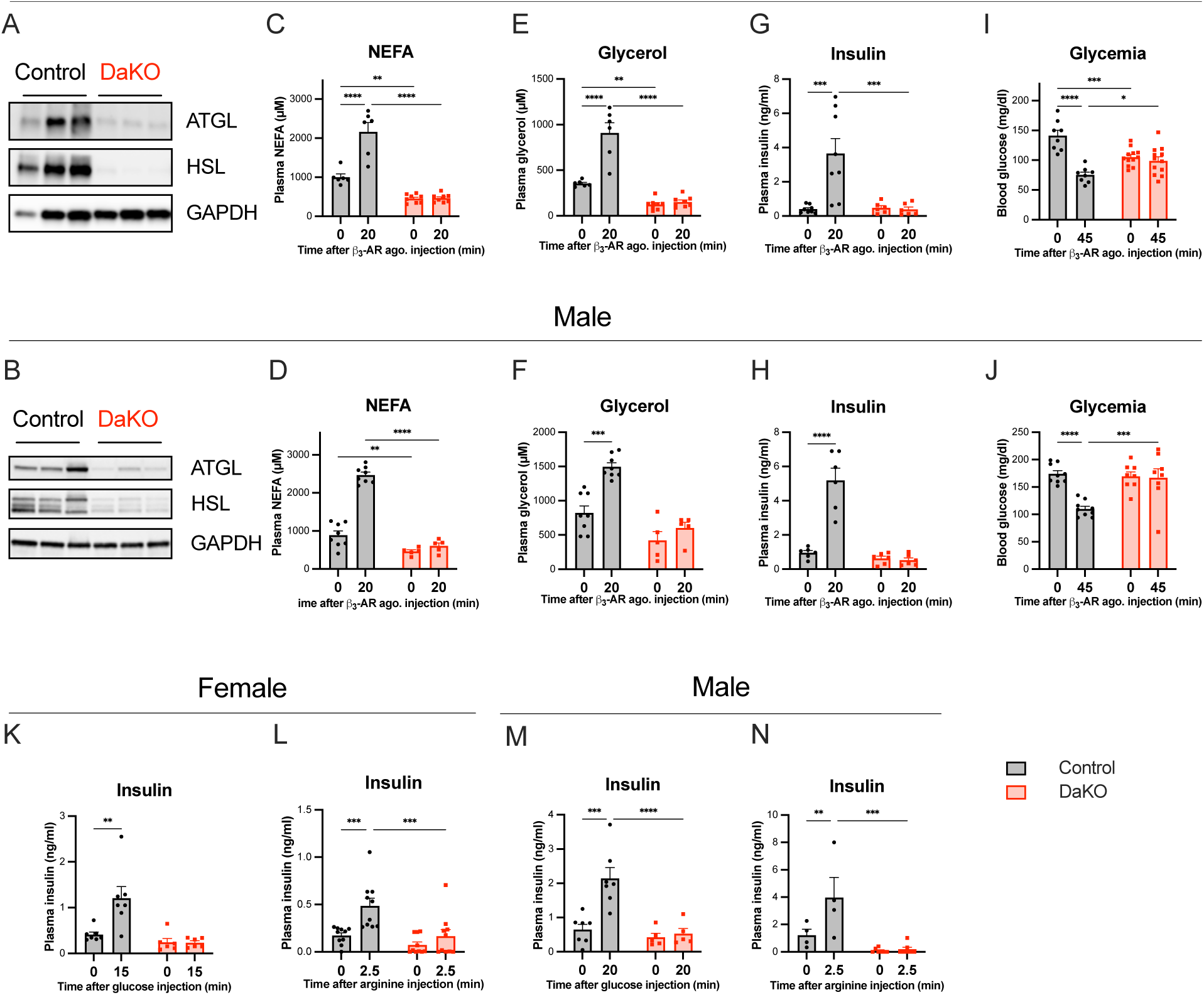
Impairment of adipocyte lipolysis affects β_3_-adrenergic receptor-, glucose- and arginine-dependent insulin secretion. (**A**-**B**) Representative protein levels of ATGL and HSL in white adipose tissues of adipocyte ATGL- and HSL-deficient mice (DaKO) and Control littermates. (**C**-**D**) Plasma non-esterified fatty acids (NEFA) (n=6-8), (**E**-**F**) plasma glycerol (n=5-8), (**G**-**H**) plasma insulin (n=5-8) and (**I**-**J**) blood glucose levels (n=8-12) in Control and DaKO mice before and after intraperitoneal injection of β_3_-adrenergic receptor (β_3_-AR) agonist (CL-316243, 1 mg/kg) at the indicated time. (**K**-**L**) Plasma insulin levels (ng/ml) in Control and DaKO female mice before and after intraperitoneal injection of (K) glucose (2 g/kg, n=6-7) or (L) arginine (1 g/kg, n=10) at the indicated time. (**M-N**) Plasma insulin levels (ng/ml) in Control and DaKO male mice before and after intraperitoneal injection of (M) glucose (2 g/kg, n=5-7) or (N) arginine (1 g/kg, n=4-7) at the indicated time. Sex is indicated, female mice (A, C, E, G, I, K and L) and male mice (B, D, F, H, J, M and N). Data are expressed as mean ± s.e.m. Statistical analyses were performed using 2-Way ANOVA with Fisher post-hoc test. *P<0.05, **P<0.01, ***P<0.001, ****P<0.0001.

To evaluate the impact of acute stimulation of adipose tissue lipolysis, control and DaKO mice of both sexes were treated with the β_3_-AR agonist CL-316243. We have previously shown that adipose tissue TG stores are rapidly mobilized after intraperitoneal injection of 1 mg/kg CL-316243^20^. Accordingly, 20 min after β_3_-AR agonist injection, we observed a drastic increase of plasma glycerol and NEFA in control mice (Figures 1C-F). Concomitantly, control mice showed a strong increase of plasma insulin (Figure 1G and 1H) accompanied by a reduction of blood glucose (Figure 1I and 1J). In male and female DaKO mice, the rise of glycerol and NEFA was completely blunted (Figures 1C-F). Strikingly, the β_3_-AR agonist had no effect on blood insulin and glucose levels in lipolysis-deficient DaKO mice (Figures 1G-J).

We next evaluated in male and female mice whether the alteration of adipocyte lipolysis influences the stimulation of insulin secretion in response to glucose and arginine. In control mice, glucose and arginine increased plasma insulin whereas no response was observed in DaKO mice (Figures 1K-N). These results suggest that the combined adipocyte ATGL and HSL deficiency blunts the stimulation of insulin secretion from pancreatic β cells *in vivo*.

### Pancreatic islets isolated from DaKO mice are functional *ex vivo*

To evaluate the functionality of DaKO Langerhans islets, we first analyzed the morphology of control and DaKO pancreatic islets. Morphometric analysis of pancreatic sections revealed comparable islet density and size (Figure 2A and 2B). Immunofluorescence analysis of pancreas sections did not reveal any difference between the two genotypes in the percentage of insulin-positive, glucagon-positive and somatostatin-positive cells (Figure 2C and 2D). Gene expression analysis of isolated islets confirmed comparable expression levels of islet cell genes, such as *Ins1*, *Gcg* and *Sst* between Control and DaKO islets of both sexes (Figure 2E and Suppl. Figure 1). Although *in vivo* insulin secretion was impaired in DaKO mice after glucose injection (Figure 1K and 1M), isolated islets from DaKO mice presented normal *ex vivo* glucose-stimulated insulin secretion (GSIS) (Figure 2F). These data show that morphology, gene expression and function of DaKO islets are preserved compared to control islets.

**Figure 2:**
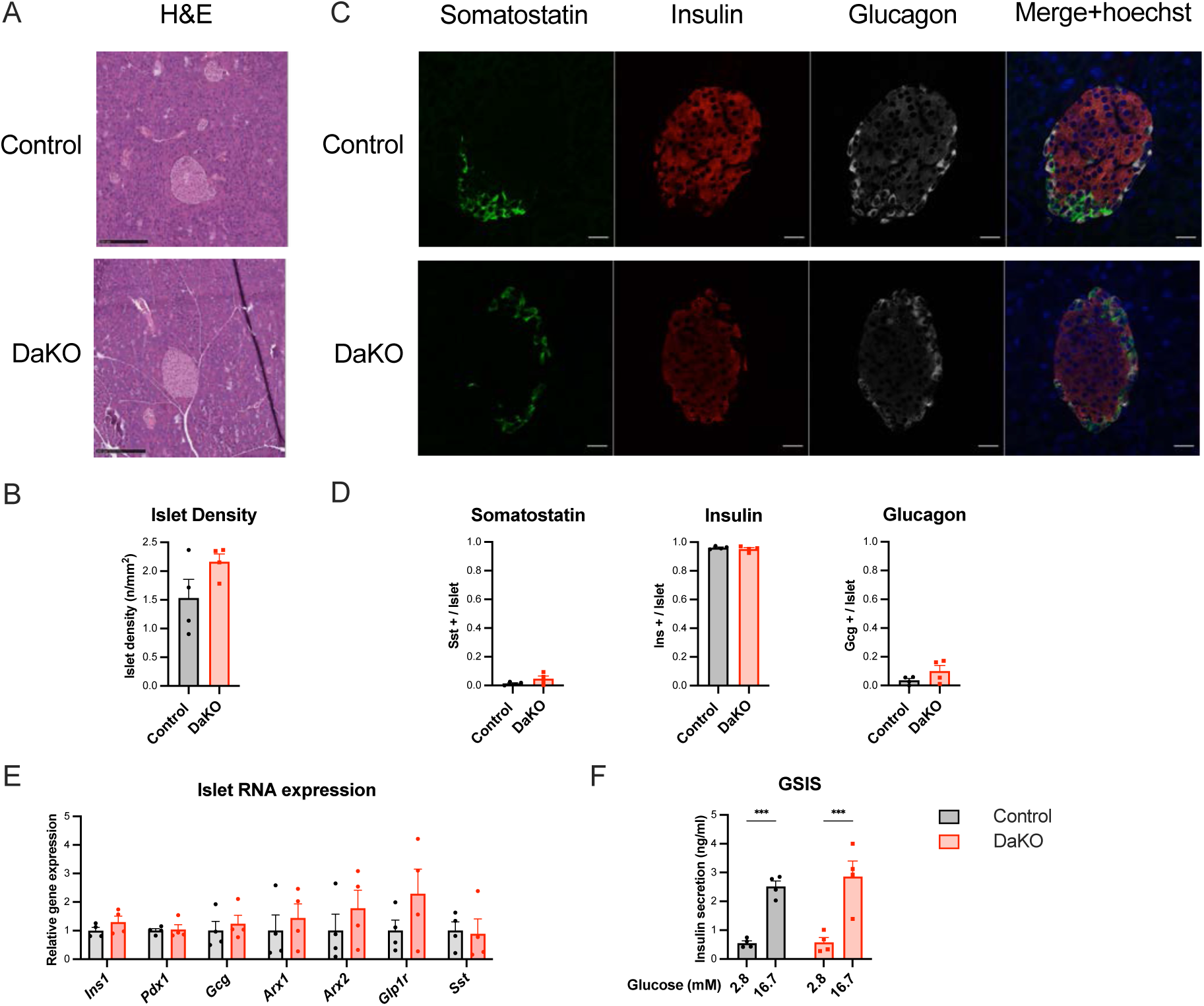
Pancreatic islets from adipocyte ATGL- and HSL-deficient (DaKO) mice exhibit normal morphology and function. (**A**) Representative pancreatic islets and (**B**) Islet density measurement from hematoxylin and eosin (H&E) staining of pancreatic sections of Control and DaKO male mice (n=4). (**C**) Representative immunofluorescence staining and (**D**) labelling quantification per islet of somatostatin (green), insulin (red), glucagon (gray) and hoechst (blue) from pancreatic sections from Control and DaKO male mice (n=4). (**E**) Relative gene expression of isolated islets from Control and DaKO male mice (n=4). (**F**) Glucose-stimulated insulin secretion (GSIS) of isolated islets from Control and DaKO female mice (n=4). Data are expressed as mean ± s.e.m. Statistical analyses were performed using unpaired t test (B-D) and ordinary two-Way ANOVA (F) with Sidak post-hoc test (E). ***P<0.001.

**Supplemental figure 1 (related to Figure 2):**
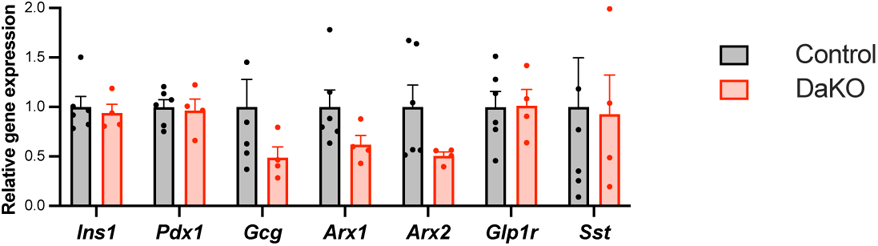
Relative gene expression of isolated islets from Control and adipocyte ATGL- and HSL-deficient (DaKO) female mice (n=4-6). Data are expressed as mean ± s.e.m. Statistical analyses were performed using ordinary two-Way ANOVA with Sidak post-hoc test.

### Ketone bodies are not implicated in adipocyte-islet interplay

As the liver is able to rapidly convert plasma NEFA to ketone bodies *via* ketogenesis, ketone body production usually follows plasma NEFA levels^21^. In addition, ketone bodies potentialize GSIS^22,23^. Opposite to control mice, blood ketone bodies were not increased in response to β_3_-AR agonist administration in DaKO mice (Figure 3A). To evaluate whether the low levels of ketone bodies could participate in the impairment of β_3_-AR-dependent insulin secretion in DaKO mice, we measured insulin secretion following β_3_-AR agonist injection in hepatocyte-specific PPARα-deficient mice (*Ppara-*HepKO) which show impairment of liver ketogenesis but intact adipose tissue lipolysis^24^. CL-316243 injection stimulated adipocyte lipolysis, as reflected by the similar increase of plasma NEFA in both control and *Ppara-*HepKO mice (Figure 3B). As expected, the deficiency in hepatic PPARα blunted the rise in blood ketone bodies following β_3_-AR agonist injection (Figure 3C). Nevertheless, the increase of plasma insulin (Figure 3D) was similar in control and *Ppara-*HepKO mice leading to a comparable reduction of blood glucose (Figure 3E). These results show that ketone bodies are not implicated in the induction of insulin secretion during lipolytic stimulation.

**Figure 3:**
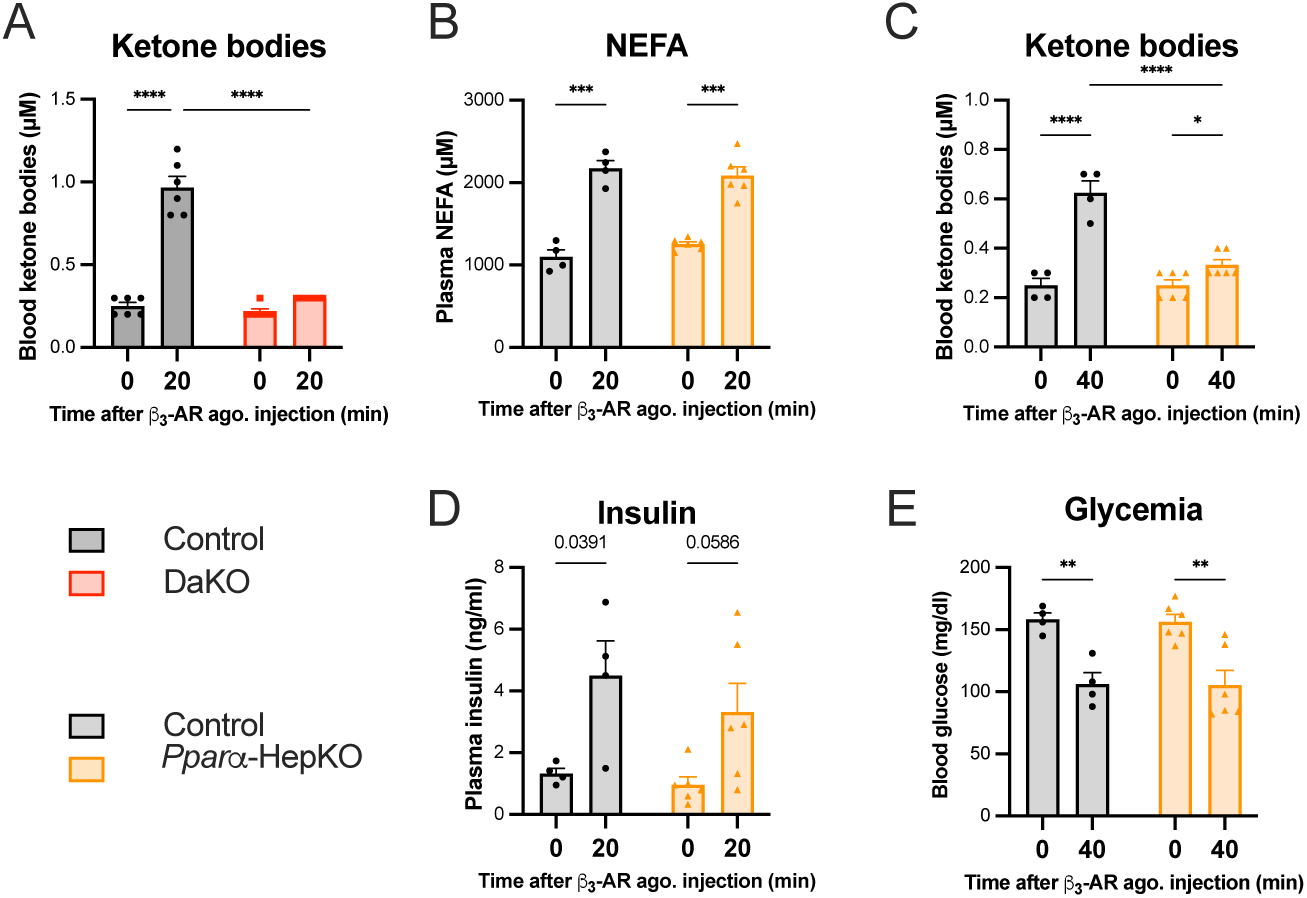
Ketone bodies are not implicated in insulin secretion following β_3_-adrenergic receptor agonist injection. (A) Blood ketone body levels of Control and adipocyte ATGL- and HSL-deficient (DaKO) male mice before and after intraperitoneal injection of β_3_-adrenergic receptor (β_3_-AR) agonist (CL-316243, 1 mg/ml) at the indicated times (n=6). (B) Plasma NEFA, (**C**) blood ketone bodies (**D**) plasma insulin and (**E**) blood glucose levels in Control and hepatocyte-specific PPARα deficient (*Ppara*-HepKO) male mice before and after intraperitoneal injection of β_3_-AR agonist (CL-316243, 1 mg/kg) at the indicated times (n=4-6). Data are expressed as mean ± s.e.m. Statistical analyses were performed using two-Way ANOVA with Sidak post-hoc test. *P<0.05, **P<0.01, ***P<0.001, ****P<0.0001.

### Adipocyte lipolysis potentiates GSIS in isolated islets

To determine whether adipocyte could directly impact islet insulin secretion, we took advantage of *ex vivo* models of primary mouse adipocytes and isolated mouse islets. We differentiated pre-adipocytes from adipose tissue stromal vascular fraction and stimulated, or not, lipolysis for 2h with β_3_-AR agonist to generate control adipocyte media or stimulated adipocyte media, containing the lipolytic products (glycerol and NEFA, Suppl. Figure 2A). This setting was used to measure the influence of this adipocyte-conditioned media on isolated mouse islets GSIS. We first verified that isolated islets were not responding to the β_3_-AR agonist (Suppl. Figure 2C). Then, we tested the capacity of control islets to secrete insulin in response to glucose in static incubations. At 1 mM glucose, the adipocyte-conditioned medium had no effect on insulin secretion (Figure 4A). As expected, 15 mM glucose stimulated insulin secretion (Figure 4A). Notably, GSIS was potentiated by the stimulated adipocyte media obtained following lipolysis stimulation but not by the control adipocyte media (Figure 4A), suggesting that adipocyte lipolysis *per se* directly modulates insulin secretion.

**Figure 4:**
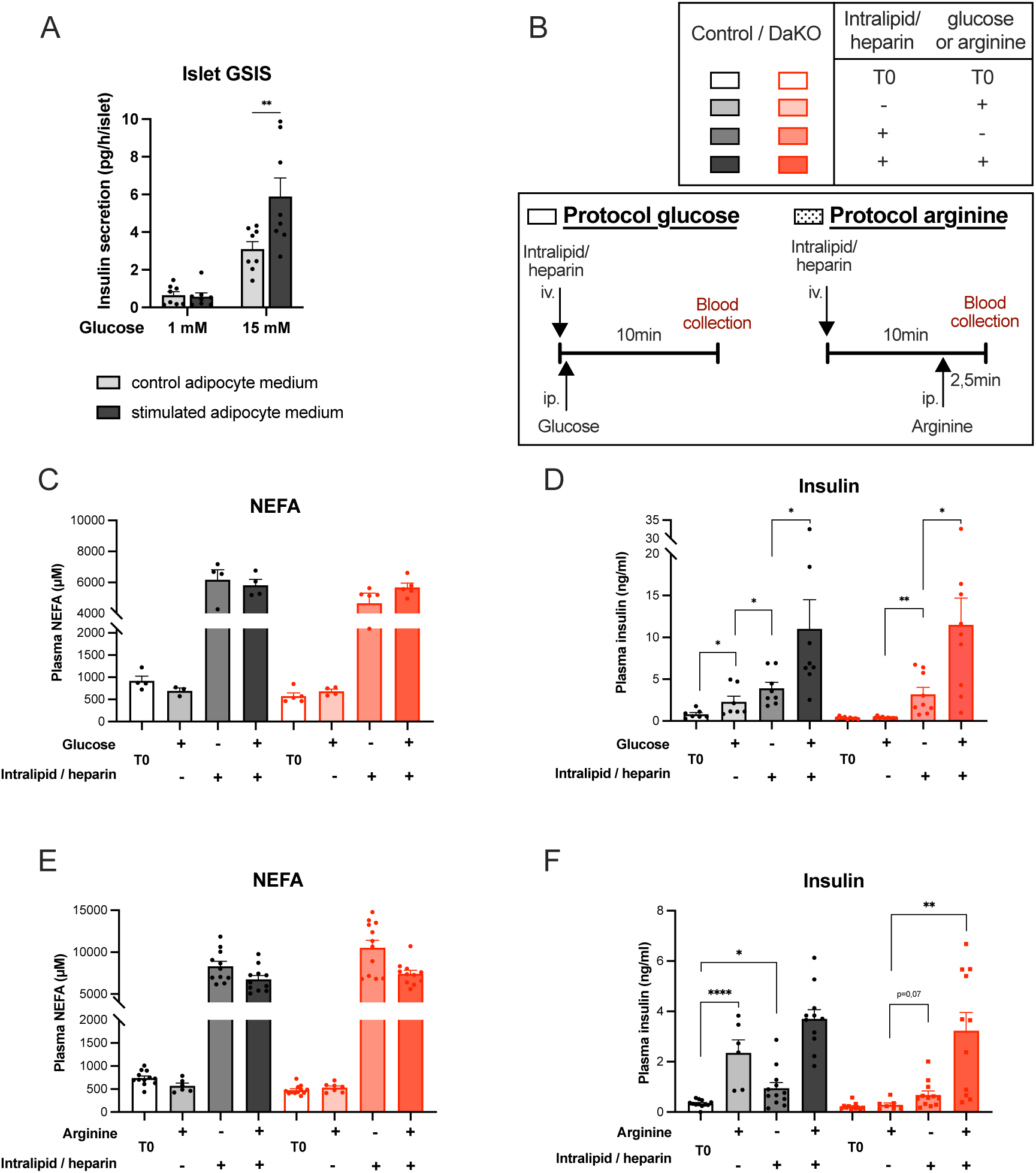
Fatty acids stimulate insulin secretion. (A) Glucose-stimulated insulin secretion (GSIS) of mouse isolated islets incubated with conditioned-medium from adipocytes or conditioned-medium from β_3_-adrenergic receptor (β_3_-AR) agonist stimulated adipocytes (n=8). Data are expressed as mean ± s.e.m. Statistical analyses were performed using two-way ANOVA with Turkey post-hoc test. ***P<0.001, ****P<0.0001. (B) *In vivo* experimental protocol performed in Control and adipocyte ATGL- and HSL-deficient (DaKO) mice to artificially mimic lipolysis using an intravenous injection of Intralipid / heparin followed by an intraperitoneal injection of glucose or arginine. (C) Plasma non-esterified fatty acids (NEFA) (n=3-5) and (**D**) plasma insulin (n=7-9) levels of Control and DaKO male mice before and after intravenous injection of 20% Intralipid / heparin (150µl) and/or intraperitoneal glucose injection (2 g/kg) as indicated. (**E**) Plasma NEFA (n=6-12) and (**F**) plasma Insulin levels (n=6-12) in Control and Dako female mice before and after intravenous injection of 20% Intralipid / heparin (150µl) and/or intraperitoneal arginine injection (1 g/kg) as indicated. Data are expressed as mean ± s.e.m. Statistical analyses were performed using paired t-test. *P<0.05, **P<0.01, ***P<0.001, ****P<0.0001.

**Supplemental figure 2 (related to Figure 4):**
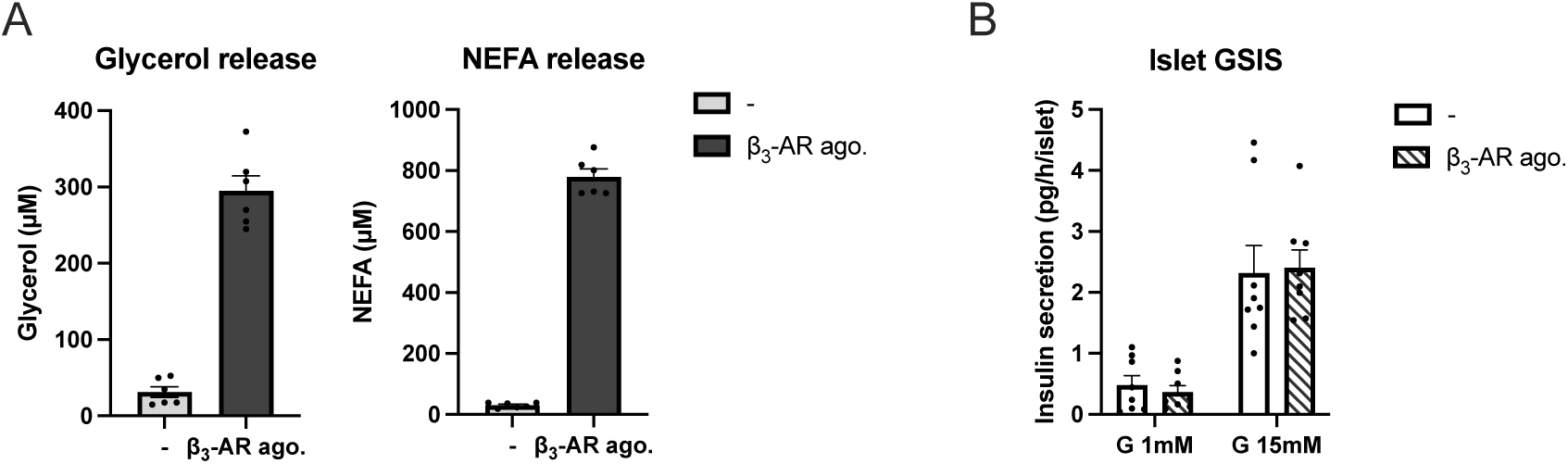
(**A**) Glycerol and non-esterified fatty acids (NEFA) in the media of differentiated adipocytes, 2 h after stimulation with the β_3_-adrenergic receptor agonist, CL-316243 100nM (β_3_-AR ago.) or not (-) (n=6). (**B**) Effect of the β_3_-AR agonist CL-316243 (100 nM) on glucose-stimulated insulin secretion (GSIS) of mouse isolated-islet (n=8). Data are expressed as mean ± s.e.m.

### An increase in plasma NEFA restores glucose- and arginine-stimulated insulin secretion in DaKO mice

To investigate the role of NEFA produced during adipose tissue lipolysis *in vivo*, we first studied the relationship between circulating NEFA and insulin levels. We observed a positive correlation between the variation of plasma NEFA levels and plasma insulin levels before and 20 min after β_3_-AR agonist stimulation (Suppl. Figure 3). To directly address the role of NEFA on insulin release *in vivo*, we performed intravenous injection of Intralipid and heparin into control and DaKO mice (Figure 4B).

**Supplemental figure 3 (related to Figure 4):**
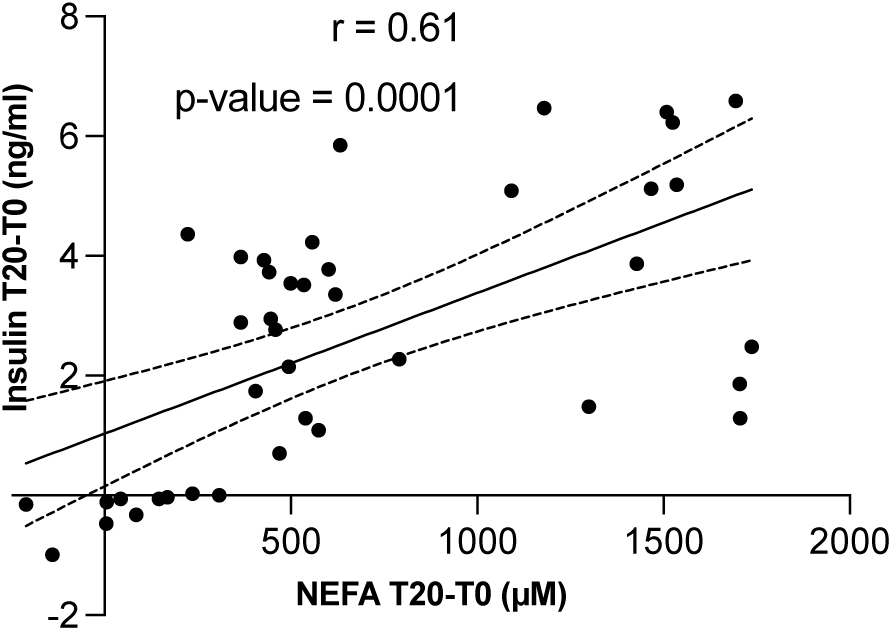
Correlation between increases in plasma insulin and NEFA (non-esterified fatty acids) of Control and DaKO mice following β_3_-AR agonist intraperitoneal administration (CL-316243, 1 mg/kg, n = 41). Pearson’s r and p-value are shown.

When Intralipid is injected with heparin, the TG emulsion is hydrolyzed into fatty acids and leads to an increase in plasma NEFA (Figure 4C). To investigate the specific contribution of NEFAs on *in vivo* GSIS, we performed the experiment concomitantly with and without intraperitoneal injection of glucose (Figure 4B). As in Figure 1K and M, glucose injection alone increased plasma insulin in control but not in DaKO mice (Figure 4D). Interestingly, injection of Intralipid and heparin without glucose increased plasma insulin to similar levels in control and DaKO mice. Moreover, when glucose was co-administered with Intralipid and heparin, GSIS was potentiated to similar levels in the two genotypes (Figure 4D). We also performed Intralipid and heparin injection with or without intraperitoneal injection of arginine (Figure 4B). Arginine had no effect on plasma NEFA levels (Figure 4E). As in Figure 1L and 1N, arginine injection alone led to an increase in insulin secretion in control but not in DaKO mice (Figure 4F). Similarly to glucose, insulin secretion induced by arginine was potentiated by Intralipid and heparin in both control and DaKO mice (Figure 4F). Therefore, our results show that mimicking lipolysis with exogenous injection of lipids restores insulin secretion in DaKO mice, and potentiates glucose- and arginine-induced insulin secretion.

### FFAR1 and FFAR4-somatostatin are involved in adipose tissue lipolysis-dependent insulin release

The fatty acid receptors FFAR1 and FFAR4 are potential mediators of adipose tissue lipolysis-dependent insulin release^6,10,25,26^. The involvement of FFAR1 is suggested by the observation that insulin secretion in response to a β_3_-AR agonist is attenuated in *Ffar1*^-/-^ mice compared to controls^27^. To establish the role of FFAR1 in our model, we administered a selective FFAR1 agonist by oral gavage before glucose injection in control and DaKO mice. The FFAR1 agonist partially rescued *in vivo* GSIS in DaKO mice (Figure 5A), suggesting that FFAR1 is involved in adipose tissue lipolysis-induced insulin secretion. Next, we investigated the FFAR4-somatostatin axis because δ-cell somatostatin is a strong paracrine inhibitor of insulin secretion, and its production is inhibited following FFAR4 activation^2,6,10,12,28^. To directly explore the potential contribution of somatostatin in lipolysis-induced insulin secretion, as we did in Figure 4A, we first tested the capacity of islets from *Sst^-/-^*mice to secrete insulin when incubated with control adipocyte-conditioned media or stimulated adipocyte-conditioned media (containing the lipolytic products). Contrary to *Sst^+/+^* islets, we did not observe GSIS potentiation by the stimulated adipocyte media in *Sst^-/-^* islets (Figure 5B). These results are in accordance with a role of δ-cell somatostatin in mediating the effects of adipocyte lipolysis on pancreatic islets. Then, to explore the upstream control of somatostatin secretion by FFAR4, we determined the induction of insulin secretion after lipolysis stimulation with β_3_-AR agonist in control and *Ffar4*^-/-^ mice. The increase in plasma NEFA and glycerol was similar in the two genotypes (Figure 5C and 5D). However, the increase in plasma insulin was blunted in *Ffar4*^-/-^ compared to control mice after β_3_-AR injection (Figure 5E). Altogether, these results suggest that the induction of pancreatic insulin secretion by adipose tissue lipolysis involves a direct effect on β cells via FFAR1 and a paracrine effect via FFAR4-somatostatin axis.

**Figure 5:**
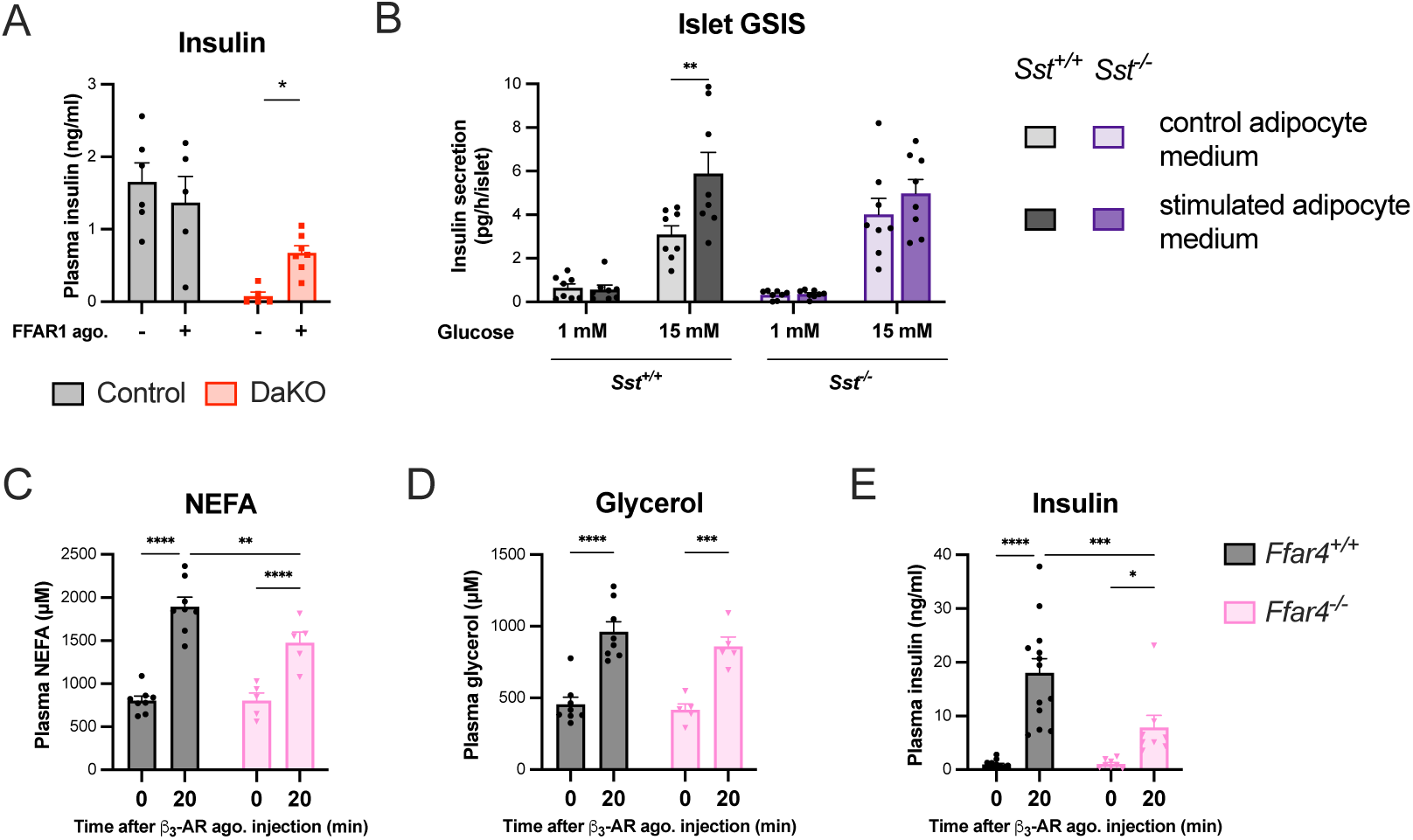
FFAR1, FFAR4 and somatostatin are involved in adipose tissue lipolysis-dependent insulin release. (A) Plasma insulin levels 15 min after intraperitoneal glucose injection (2 g/kg) in Control and adipocyte ATGL- and HSL-deficient (DaKO) female mice (n=5-7). FFAR1 agonist (1 mg/kg) or vehicle was administrated by gavage 30 min before glucose injection. Data are expressed as mean ± s.e.m. Statistical analyses were performed using Kolmogorov-Smirnov nonparametric unpaired test to test the effect of FFAR1 agonist in each genotype. *P<0.05. (B) Glucose-stimulated insulin secretion (GSIS) of mouse isolated islets from *Sst^+/+^* and Sst*^-/-^* mice incubated with conditioned-medium from adipocytes or conditioned-medium from β_3_-AR agonist stimulated adipocytes (n=8). Data are expressed as mean ± s.e.m. Statistical tests between adipocyte media treatment were performed using two-way ANOVA with Sidak post-hoc test. **P<0.01. Control *Sst^+/+^* control data were presented in Figure 4A (C) Plasma NEFA (n=5-8), (**D)** plasma glycerol (n=5-8) and (**E**) plasma insulin (n=8-13) levels in *Ffar4^+/+^* and *Ffar4^-/-^*male mice following intraperitoneal β_3_-AR agonist injection (CL-316243, 1 mg/kg) at the indicated time. Data are expressed as mean ± s.e.m. Statistical analyses were performed using 2-Way ANOVA with Sidak post-hoc test. *P<0.05, **P<0.01, ***P<0.001, ****P<0.0001

### Adipocyte lipolysis stimulates insulin secretion in mice with diet-induced obesity

We investigated whether the control of insulin secretion by adipose lipolysis is preserved in the pathological context of diet-induced obesity. To this end, we challenged Control and DaKO mice for 12 weeks with a high-fat diet (HFD). In a first protocol, deficiency in ATGL and HSL was induced before the start of HFD (Figure 6A). In a second protocol, deficiency in the neutral lipases was induced once diet-induced obesity was installed. A difference in body weight was observed between the two genotypes in the first but not the second protocol (Figure 6B and 6C). At approximately 10 weeks of HFD, mice were subjected to lipolysis stimulation. In both protocols, the stimulation of lipolysis by the β_3_-AR agonist increased plasma NEFA levels in Control mice (Figure 6D and 6E). Comparatively, this increase was markedly blunted in DaKO mice (Figure 6D and 6E). In both protocols, lipolysis induction of plasma insulin was largely abrogated in DaKO mice (Figure 6F and 6G). Accordingly, we observed a reduction of blood glucose in Control mice but not in DaKO mice (Figure 6H and 6I).

**Figure 6:**
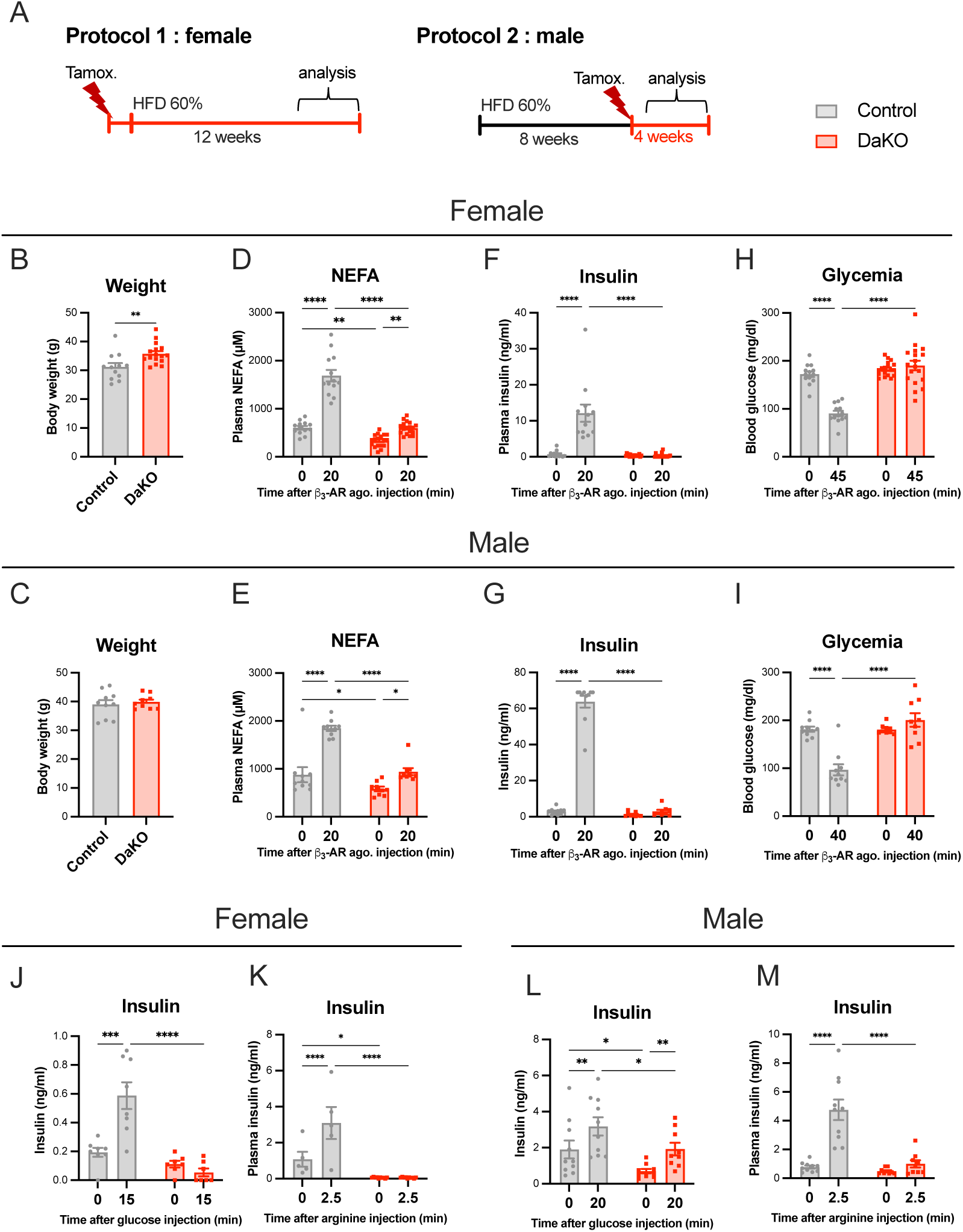
Adipocyte lipolysis stimulates insulin secretion in diet-induced obesity mouse model. (**A**) General description of the high fat diet (HFD) protocols used in this study. (**B-C**) Body weight, (**D-E**) Plasma non-esterified fatty acids (NEFA), (**F-G**) plasma insulin and (**H-I**) blood glucose levels in Control and DaKO HFD-mice before and after intraperitoneal injection of β_3_-adrenergic receptor (β_3_-AR) agonist (CL-316243, 1 mg/kg) at the indicated time (n=13-18 females; n=9-10 males). (**J-K**) Plasma insulin levels (ng/ml) in Control and DaKO HFD-female mice before and after intraperitoneal injection of (**J**) glucose (2 g/kg, n=7-8) or (**K**) arginine (1 g/kg, n=5-10) at the indicated time. (**L-M**) Plasma insulin levels (ng/ml) in Control and DaKO HFD-male mice before and after intraperitoneal injection of (**L**) glucose (2 g/kg) or (**M**) arginine (1 g/kg) at the indicated time (n=9-10). Sex is indicated, female mice (B, D, F, H, J and K), male mice (C, E, G, I, L and M). Data are expressed as mean ± s.e.m. Statistical analyses were performed using 2-Way ANOVA with Sidak post-hoc test. For body weight, statistical analyses were performed using Kolmogorov-Smirnov nonparametric unpaired test to compare genotype. *P<0.05, **P<0.01, ***P<0.001, ****P<0.0001.

To further confirm the importance of this mechanism in HFD, we next assessed whether glucose- or arginine-dependent insulin secretion was also impacted by the defect of adipose tissue lipolysis, as shown in chow-fed mice (Figure 1K and 1M). In response to glucose injection, the increase in plasma insulin of HFD-fed Control mice was not observed or reduced in HFD-fed DaKO mice (Figure 6J). This inability of HFD-fed DaKO mice to stimulate insulin secretion was also seen during arginine injection (Figure 6K).

As in chow-fed mice (Figure 2), we checked that there was no alteration of pancreas morphology and function in HFD-fed DaKO mice compared to HFD-fed Control mice. Control and DaKO islets exhibit normal morphology and function during HFD. Morphometric analysis of pancreatic sections revealed comparable islet density and insulin-positive, glucagon-positive and somatostatin-positive cells between Control and DaKO islets (Suppl. Figures 4A-D). Pancreatic islet insulin content as well as GSIS were similar between Control and DaKO mice (Suppl. Figure 4E and 4F).

Together, our results indicate that the regulation of insulin secretion by adipose tissue lipolysis is preserved in the context of diet-induced obesity.

**Supplemental figure 4 (related to Figure 6):**
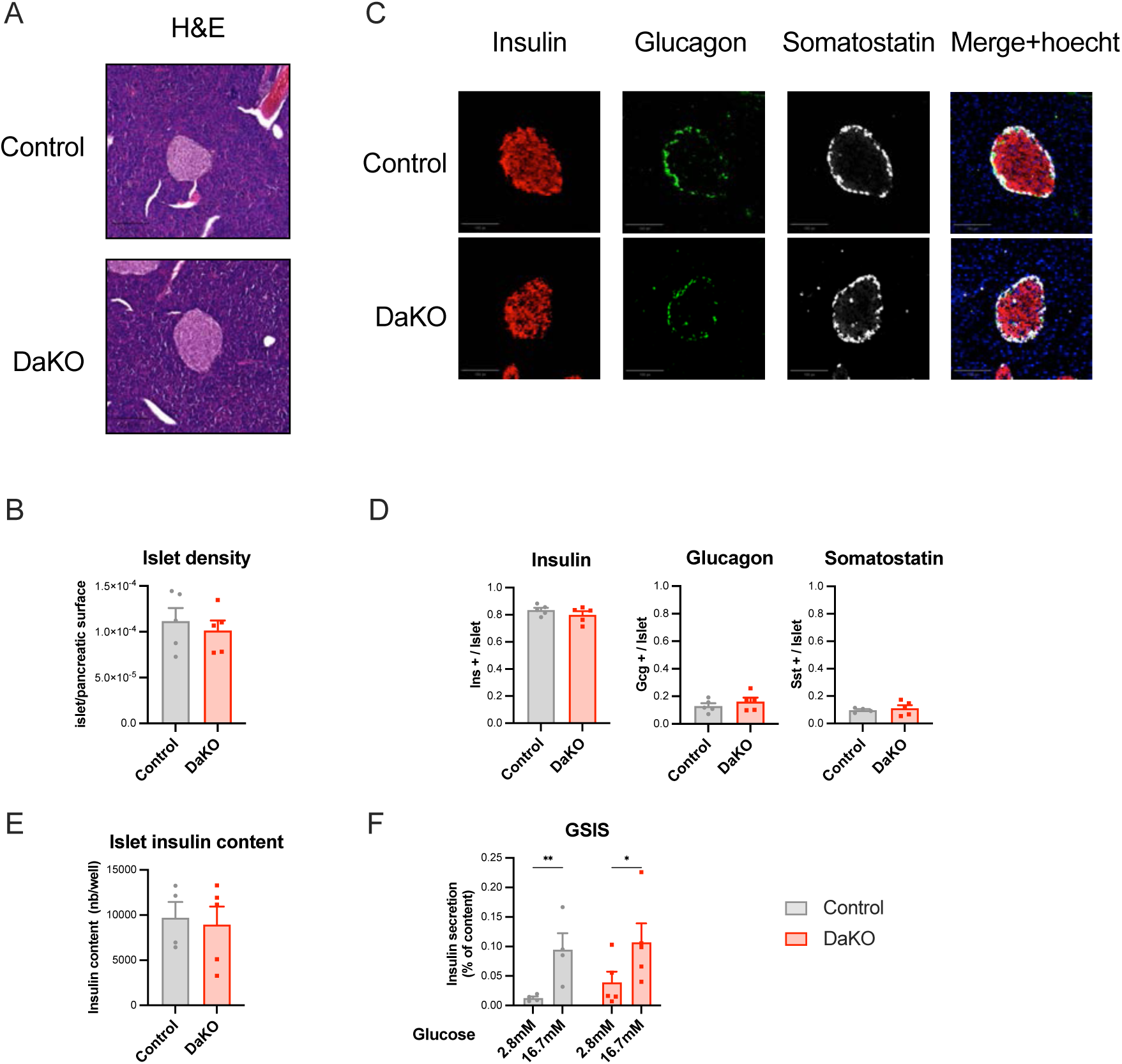
Pancreatic islets from high fat diet-fed (HFD) adipocyte ATGL- and HSL-deficient mice (DaKO) exhibit normal morphology and function compare to islets from Control mice. (A) Representative hematoxylin and eosin (H&E) staining of islet from pancreatic sections of Control and DaKO male mice submitted to HFD (n=5). (B) Islet density measurement from immunofluorescence staining of insulin and glucagon pancreatic sections from Control and DaKO male mice submitted to HFD (n=5). (C) Representative immunofluorescence staining of somatostatin, insulin, glucagon and hoechst in pancreatic sections from Control and DaKO male mice submitted to HFD (n=5). (D) Insulin, glucagon and somatostatin labelling quantification per islet from Control and DaKO mice submitted to HFD (n=4-5). (E) Insulin content of isolated islets from Control and DaKO female mice submitted to HFD (n=3-5). (F) Glucose-stimulated insulin secretion (GSIS) in isolated islets from Control and DaKO female mice submitted to HFD (n=3-5). Data are expressed as mean ± s.e.m. Statistical analyses were performed using Kolmogorov-Smirnov nonparametric unpaired test to compare genotype. For GSIS, statistical analyses were performed using two-Way ANOVA with Fisher post-hoc test. **P<0.05, **P<0.01.

## DISCUSSION

Early studies suggest that free fatty acids, unlike triglycerides, enhance glucose-stimulated insulin secretion^4,19,29–32^. However, the specific contribution of adipose tissue lipolysis, the primary source of circulating fatty acids, has not been thoroughly investigated^25,27,33^. This may be explained by the fact that, at first glance, it seems counterintuitive that an energetic substrate produced during the fasting state would stimulate insulin release. The results presented here provide evidence that the genetic ablation of adipocyte lipolysis in mice leads to defective insulin secretion in response to administration of a β_3_-AR agonist, as well as to glucose or arginine injection. The lack of morphological or functional defects in the pancreatic islets strongly indicates direct inter-organ interplay. We demonstrate that the products of adipose tissue lipolysis, NEFAs, rather than ketone bodies play a key role in this process. Indeed, an artificial increase of plasma NEFA levels using Intralipid and heparin restores insulin secretion in DaKO mice. Moreover, our findings illuminate that the FFAR1 and FFAR4-somatostatin pathways both contribute to the mechanism of lipolysis-dependent insulin release. Interestingly, we provide conclusive evidence that the acute control of insulin secretion by adipocyte lipolysis is conserved in the pathological mouse model of obesity induced by HFD.

Glucose is the primary regulator of insulin secretion. While the impact of fatty acids on insulin secretion is well demonstrated and documented^34^, the link between adipose tissue lipolysis and insulin secretion is not conceptually obvious^25^. During feeding, increased insulin levels due to increased glycemia inhibit adipose lipolysis, which remains too low to impact insulin secretion in return^32,35^. Conversely, during fasting, insulin levels are low and catecholamines stimulate adipocyte lipolysis which results in a rise of plasma NEFAs^2,3,13^. This complicates the evaluation of this interaction for determining whether the increase in fatty acids release from adipocyte lipolysis modulates insulin secretion. By using mice with impaired adipocyte lipolysis, our study provides evidence for this link.

We hypothesize that adipocyte lipolysis could modulate insulin secretion during nutritional transitions. The preparation of the body for feeding, including sensory perception of food (sight, smell, taste)^36^, may provide a link between the central nervous system, adipose tissue lipolysis and pancreatic insulin secretion. A recent study reveals that food perception (smell) activates the sympathetic nervous system during fasting, and increases plasma NEFAs levels through stimulation of adipose tissue lipolysis^37^. Additionally, it is well documented that food perception stimulates insulin release in mice via the cephalic phase insulin response. This occurs prior to nutrient absorption and is physiologically important because it enables the body to adapt to food arrival, minimizing postprandial hyperglycemia^38,39^. Therefore, based on the data presented, adipose tissue lipolysis could be involved in the integration of the cephalic insulin response; the sympathetic nervous system activation by food perception inducing adipose tissue lipolysis which stimulates insulin secretion.

During obesity, adipocyte dysfunction is observed^13,40,41^. While catecholamine resistance occurs during HFD^42–44^, the rise in plasma NEFA level remains substantial. Our work shows that the acute control of insulin secretion by adipocyte lipolysis is conserved and sufficient to induce insulin secretion in a mouse model of diet-induced obesity. This is not the case when adipocyte lipolysis is abrogated. Our findings suggest that, in the context of obesity, the adipose tissue lipolysis induction of insulin secretion could contribute to hyperinsulinemia. Currently, it is not entirely clear what is causing hyperinsulinemia^45^. Interestingly, recent studies have highlighted that hyperinsulinemia may be a consequence of NEFA elevation, which would actually constitute an early alteration driving insulin resistance in obesity^14–16,46^. Our results support this pathophysiological model positioning islet overstimulation due to adipose tissue lipolysis NEFAs as an early defect.

Mechanistically, we identified potential pathways explaining the action of adipose tissue NEFAs release on insulin secretion: 1-the activation of β-cell FFAR1 and, 2-the activation of δ-cell FFAR4 and the somatostatin paracrine pathway. First, the data in DaKO mice treated with a FFAR1 agonist are consistent with those reported in *Ffar1*^-/-^ mice, in which insulin secretion induced by adipocyte lipolysis stimulation is reduced by approximately 40%^27^. Second, regarding FFAR4 and somatostatin, we show that the absence of adipocyte lipolysis following β_3_-AR agonist injection blunts insulin secretion. Recent findings have suggested that δ-cell FFAR4 controls somatostatin secretion, ultimately regulating β-cell hormonal secretion^10^. Our results in *Sst^-/-^* islets and *Ffar4^-/-^* identify the FFAR4-somatostatin pathway as a partial mediator of insulin secretion by adipocyte lipolysis.

In summary, our findings demonstrate that adipose tissue lipolysis plays a critical role in the regulation of insulin secretion and indicate that multiple metabolic and signaling pathways mediate the effects of NEFAs on pancreatic islets. The magnitude of the lipolysis-induced modulation of insulin secretion underscores the need for further investigations to define the temporal dynamics of this regulatory mechanism under physiological and pathological conditions.

## Limitation of the study

The timing of the protocol that we used for arginine-induced insulin secretion (2.5 min) suggests a rapid crosstalk between adipocytes and islets. Nevertheless, we cannot rule out the possibility of intermediary factors other than fatty acids being involved in the process of adipocyte lipolysis and β-cell insulin secretion. Indeed, it is possible that other factors, or combinations of factors (e.g. NEFAs and adipocyte lipocalins), might play a role in the mechanism. In our study, our findings exclude ketone bodies and largely support the role of NEFAs, as these restore insulin secretion in DaKO mice. To date, as reported in the literature, there is no evidence supporting that glycerol plays a role in the process^47,48^. In our study, we were not able to discriminate the autocrine activation of FFAR1 by fatty acids and other lipid derivatives released locally by ß cells during glucose stimulation^7,8^. Nevertheless, our findings support the role of FFAR1 activation by lipolytic products. Finally, investigating the role of adipocyte lipolysis in the control of glucagon secretion is of interest and will require further investigations.

## Acknowledgements

We are grateful to members of the Langin-Gourdy Team (I2MC, Toulouse) for useful discussion and feedback. Geneviève Tavernier (I2MC) managed transgenic mouses lines. Anne Fougerat (Toxalim, Toulouse), Khalil Acheikh-Ibn-Oumar (I2MC), Emma Novarini (I2MC) and Fernando Cortes-Camacho (I2MC) provided technical assistance. We thank Hervé Guillou, Walter Walli and Alexandra Montagner for the gift of *Ppara-*HepKO mice^24^. AstraZeneca (Gothenburg, Sweden) generously provided chemical compounds. The following technological facilities contributed to the work: GenoToul Anexplo (CREFRE, Toulouse), GenoToul GeT Santé (Emeline Lhuillier, I2MC, Toulouse), GenoToul TRI (Christèle Ségura, I2MC). This study was supported by Inserm (to PDD), European Research Council (ERC) under the European Union’s Horizon 2020 research and innovation program (SPHERES, ERC Synergy Grant agreement no. 856404 to DL), European Foundation for the Study of Diabetes (EFSD-Boehringer Ingelheim European Research Programme on “Multi-System Challenges in Diabetes” 2024 to PDD and MR), Agence National de la Recherche (ANR-25-CE14-7667-02 to PDD and MR; ANR-20-CE16-0024-03 to JSA; ANR-23-CE14-0040-01 to JSA; ANR-25-CE14-6544-02 to JSA), Université de Lille and Initiative d’Excellence de l’Université de Lille (CDP Mosaic to JSA), Institut Pasteur de Lille (CTRL Melodie to JSA), Fondation pour la Recherche Médicale (EQU202103012732 to JSA), Société Francophone du Diabète (SFD23 to EC and SFD25-KaT2D to JSA).

## Author contributions

CFF, DL and PDD conceptualized the work. CFF, MM, SCS, EM and PDD performed the experiments using DaKO mice, *Ppara-*HepKO mice and primary adipocytes, and analyzed the data. ET, AD and PG performed and supervised GSIS experiment on pancreatic islet from *Sst^+/+^* and *Sst^-/-^* with adipocyte-conditioned media. LR and VP performed and supervised experiment on *Ffar4*^-/-^ mice. AP and MR performed and supervised GSIS experiments on pancreatic islets isolated from HFD-fed mice. EC, LR and JSA performed and supervised pancreatic islet morphology and GSIS experiments, pancreas and islet analysis. CFF and PDD analyzed all the data. DL and PDD supervised the project and wrote the manuscript. All the authors reviewed and edited the manuscript.

## MATERIAL AND METHODS

### Key resources tab

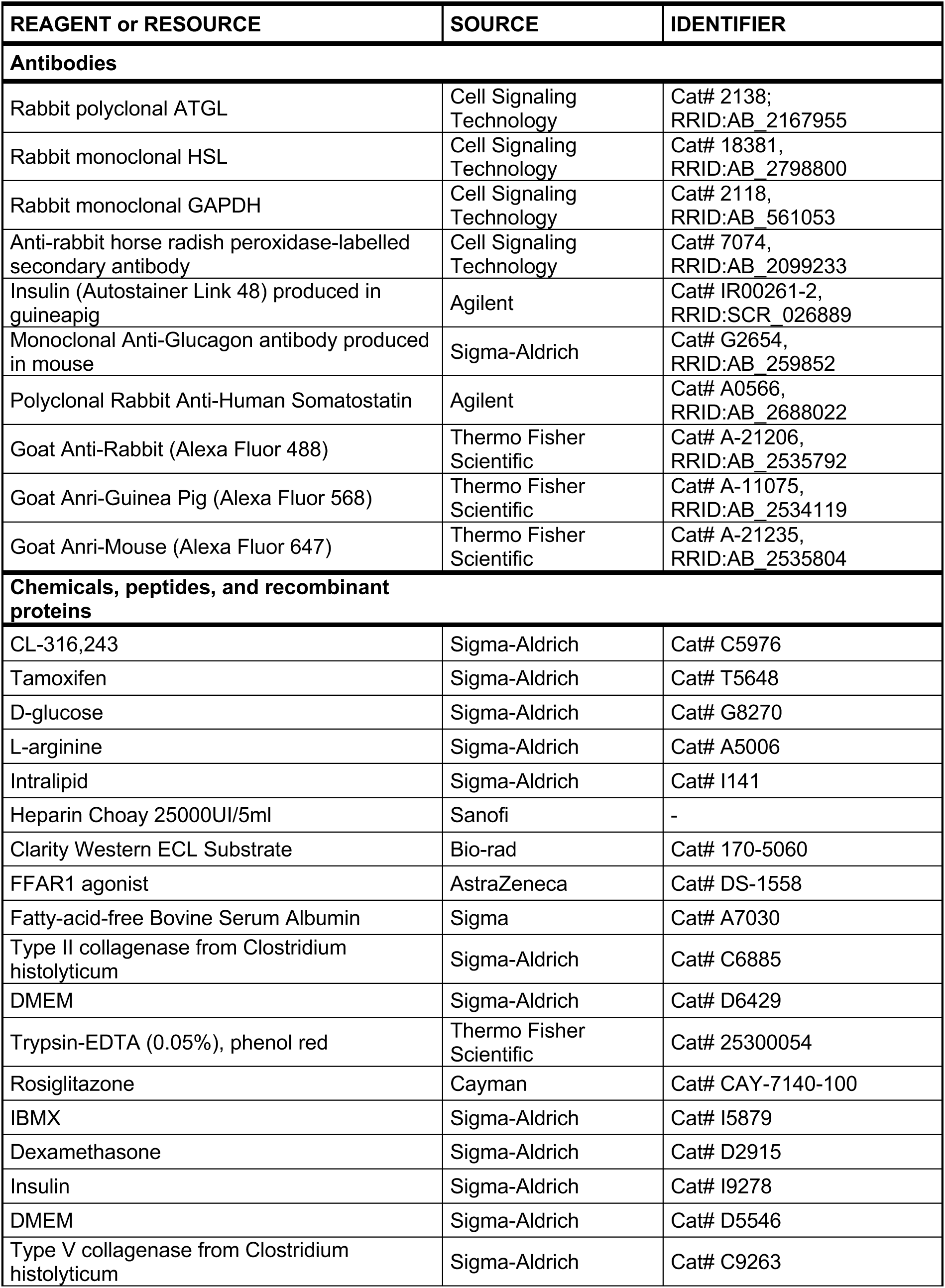
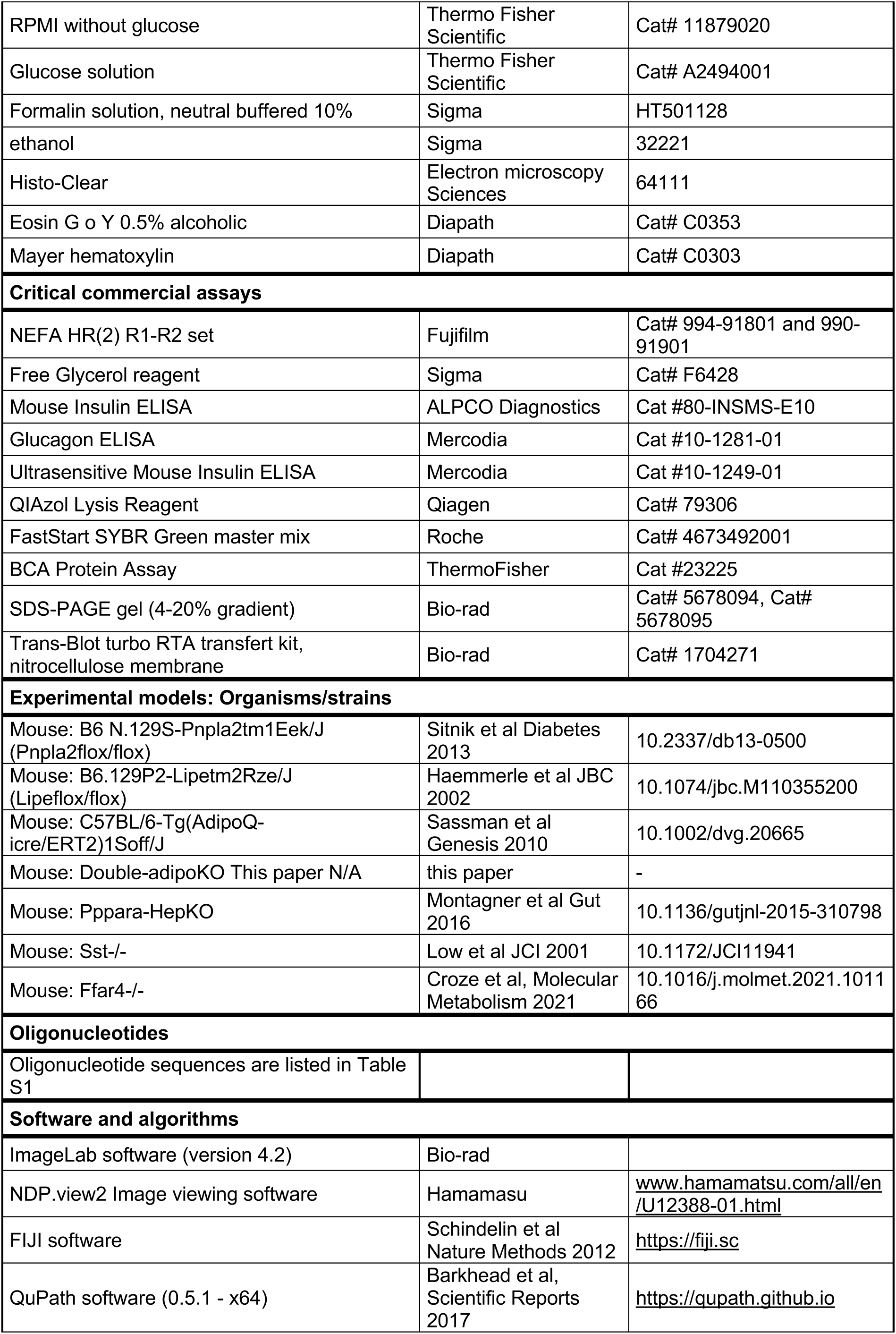
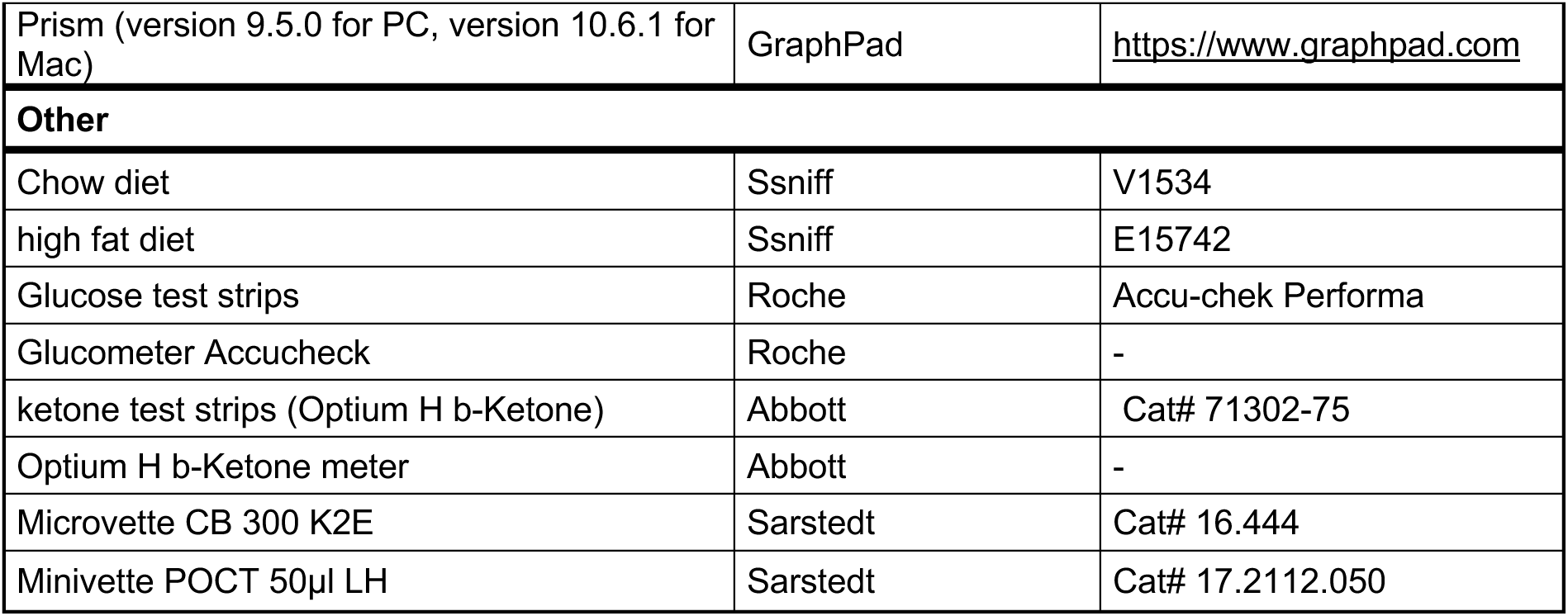

### Mouse models

We generated a tamoxifen-inducible mouse model deficient in adipocytes for both *Pnpla2* (encoding ATGL) and *Lipe* (encoding HSL) by crossing *Pnpla2^fl/fl^* mice^49^, *Lipe^fl/fl^* mice^50^ and Tg(AdipoQ-Cre/ERT2)1Soff/J transgenic mice^51^. Tg(AdipoQ-Cre/ERT2)1Soff/J mice express the tamoxifen-inducible Cre/ERT2 recombinase, under the control of the adipocyte-specific adiponectin promoter. Around 7 to 10 weeks of age, triple transgenic mice received tamoxifen (1 mg/day by oral gavage for 5 days, Sigma) to induce adipocyte deletion of ATGL and HSL. Same sex littermates, *Pnpla2 ^fl/fl^*; *Lipe^fl/fl^* ;AdipoQ-Cre/ERT2^-^ mice treated with tamoxifen were used as control. The hepatocyte-specific *Ppar*α knockout mouse (*Ppara-*HepKO), the somatostatin knockout mouse (*Sst*^-/-^) and the *Ffar4* knockout mouse (*Ffar4*^-/-^) lines were previously described^10,24,52^.

Mice were fed standard chow (V1534 Ssniff) and housed according to Inserm guidelines and European Directive 2010/63/UE in the local animal care facility (agreements A 31 555 04 and C 31 555 07). Protocols were approved by the French Ministry of Research (APAFIS #34176-2021112915333477) after review by local ethical committee (Comité d’éthique en expérimentation animale de l’UMS006/CREFRE, CEEA122, Toulouse, France).

### β_3_-AR agonist injection

Around 9:00 a.m, 1 mg/kg body weight of CL-316243 (Sigma) was administrated intraperitoneally (i.p.) to 1h fasted mice. Blood and plasma parameters were determined as indicated below. Blood was collected before and 20 min after CL-316243 injection, for plasma analysis. Glycemia was measured before and after CL-316243 administration at the indicated times in the legends.

### Glucose, arginine injections and FFAR1 agonist administrations

After 4-5 h fasting, 2 mg/g body weight of glucose (Sigma) or 1 mg/g body weight arginine (Sigma) was administered by i.p. Blood was collected before and at the time indicated after injection for plasma analysis. The FFAR1 selective agonist, DS-1558 (Daiichi Sankyo), was administrated 30 min before glucose i.p. by oral gavage.

### Intralipid and heparin injection with glucose or arginine

Around 11:00 a.m, 150 µL of 20% Intralipid (Sigma) with heparin at 50 U (Heparin Choay) were administrated intravenously (i.v.) in 1h fasted mice. Immediately after, 2 mg/g body weight of glucose (Sigma) was administrated by i.p. Blood was collected before and 10 min after Intralipid injection. Same protocol was used for arginine injection, except that 1 mg/g body weight of arginine was administrated by i.p. 7,5 min after Intralipid and heparin. Blood was collected before and 10 min after of Intralipid injection (2.5 min after the arginine injection).

### High-fat diet

Two different protocols were used. Female mice were fed a high-fat diet (HFD) containing 60% fat (Ssniff E15742) for 12 weeks. Tamoxifen-induced deletion of ATGL and HSL was performed one week prior to the start of the HFD. Experimental procedures were initiated 8 weeks after start of HFD. For male mice, the same HFD was administered, except that the adipocyte deletion of ATGL and HSL induced after 8 weeks on HFD, when obesity was already installed.

### Plasma and blood analyses

Blood glucose and ketone bodies levels were measured using a glucometer (Accucheck, Roche) and a ketone body meter (Freestyle Optium H meter, Abbott), respectively. For plasma analysis, blood was collected through the tail vein at the indicated times or after sacrifice using EDTA-coated microvettes or minivettes (Sarstedt). Plasma NEFA and glycerol were measured using enzymatic assay (Fujifilm) and Free Glycerol reagent (Sigma). Plasma insulin and glucagon were measured using ultrasensitive ELISA immunoassays (ALPCO and Mercodia, respectively).

### Pancreatic islet histology

Freshly collected tissue was fixed in 10% formalin (Sigma) for 24h at room temperature and dehydrated using ethanol and histoclear II (Electron microscopy Sciences). Following paraffin embedding, blocks were cut in 4 μm slices. Following hematoxylin-eosin staining, images were obtained with nanozoomer (Hamamatsu) or Axioscan7 (Zeiss) and visualized using NDP view (Hamamatsu) and QuPath software. Immunofluorescence and immunohistochemistry were performed as described previously^53^. Pancreatic tissues were fixed as describe above. For immunofluorescence microscopy analyses, after antigen retrieval using citrate buffer, 5-μm formalin-fixed paraffin embedded (FFPE) pancreatic sections were incubated with the indicated antibodies: guinea pig anti-insulin (IR00261-2, Agilent), mouse anti-glucagon (G2654, Sigma) and rabbit anti-somatostatin (A056601, Agilent). Immunofluorescence staining was revealed by using Alexa-conjugated secondary antibody (anti-guinea pig alexa fluor 568, anti-mouse alexa fluor 647, anti-rabbit alexa fluor 488). Nuclei were stained with Hoechst. For morphometric analysis, four animals from each genotype were analyzed. Images were obtained with Axioscan (Zeiss), processed and quantified using FIJI software by an observer blinded to experimental groups. Images were visualized FIJI and QuPath software.

### Pancreatic islet studies

Mouse islets were isolated following enzymatic digestion of the pancreas with type V collagenase (1.5 mg/ml, C9263 sigma) for 10 min at 37°C as described previously^53^. Briefly, after digestion and separation in a density gradient medium, islets were purified by handpicking under a microscope and cultured during 16 h before subsequent analysis. For glucose-stimulated insulin secretion (GSIS) tests, approximately twenty islets were exposed for 1h to indicated glucose concentrations in Krebs-Ringer bicarbonate HEPES buffer containing 0.5% fatty-acid-free Bovine Serum Albumin (Sigma). Insulin released in the medium was quantified by ELISA or radioimmunoassay (80-INSMS-E01 ALPCO, 10-1113-01 Mercodia or homemade radioimmunoassay assay). For expression studies, isolated mouse islets were snap-frozen in liquid nitrogen before RNA extraction. Total RNA was extracted using Quiazol reagent (Qiagen). mRNA expression was measured after reverse transcription by quantitative RT-PCR (qRT-PCR) with FastStart SYBR Green master mix (Roche) using a LightCycler Nano or LC480 instrument (Roche). qRT-PCR results for *Ins1*, *Pdx1*, *Gcg*, *Arx1*, *Arx2*, *Glp1r* and *Sst* were normalized to endogenous *Ppia* reference mRNA levels. Results are expressed as the relative mRNA level of a specific gene expression using the formula 2^-ΔCt^.

### Primary culture of mouse adipocytes

Adipocyte progenitors are isolated from adipose tissue of 1- to 3-day-old mice, as previously described^54^. Briefly, subcutaneous fat pads are harvested, minced, and digested with type II collagenase (1 mg/mL, C688 Sigma) in PBS + 2% BSA, (Sigma) approx. 15 minutes at 37°C. After centrifugation 10 min at 500 rpm, cells were wash and plated in DMEM (Sigma) with 10% serum. The next day, the medium was changed. At approximately 70–80% confluence (3–4 days after seeding), the cells are trypsinized (ThermoFisher) and then reseeded at a density of 80,000 cells per 6-well plate well (TPP) to reach confluence 3 days later. Adipocyte differentiation is then induced using a differentiation cocktail for 2 days: Rosiglitazone (170 nM, Cayman), IBMX (500 µM, Sigma), dexamethasone (1 µM, Sigma), and insulin (170 nM, Sigma) in DMEM medium containing 1 mM glucose (Sigma) supplemented with 10% SVF. The cells are then cultured in DMEM 10% SVF, and 10 mM insulin. After 7 days of differentiation, the adipocytes were deprived of serum and insulin. The following day, adipocyte lipolytic activity is stimulated 2 hours by the β_3_-AR agonist (100 nM CL-316 243, Sigma) in RPMI medium (ThermoFisher) supplemented at 1 mM glucose. The medium was harvested after 2 hours of stimulation.

### Immunoblotting

Tissues were homogenized in lysis buffer (50 mM Tris pH8, 150 mM NaCl, 5 mM EDTA, 1% Triton) containing protease and phosphatase inhibitors (Sigma) using Precellys evolution homogenizer (Bertin Scientific) and centrifuged. Supernatants were harvested for determination of total protein concentration. Equal amounts of solubilized proteins were loaded on 4-20% gradient SDS-PAGE gels (Bio-Rad), blotted onto nitrocellulose membranes and incubated overnight with primary antibodies, rabbit polyclonal anti-ATGL (Cell Signaling Technology); rabbit monoclonal anti-HSL (Cell Signaling Technology) and rabbit monoclonal anti GAPDH (Cell Signaling Technology). Subsequently, immuno-reactive proteins were blotted with anti-rabbit horseradish peroxidase-labelled secondary antibodies for 1h at room temperature and revealed by enhanced chemiluminescence reagent (Bio-Rad), visualized using ChemiDoc Touch Imaging System and data analyzed using the ImageLab 4.2 version software (Bio-Rad).

### Statistical Analysis

Data are presented as mean ± SEM. Statistical analysis was performed using Prism 10 software (GraphPad). Tests are indicated in the legends. Differences were considered statistically significant at p<0.05.

## Supplemental information

**Table S1.**
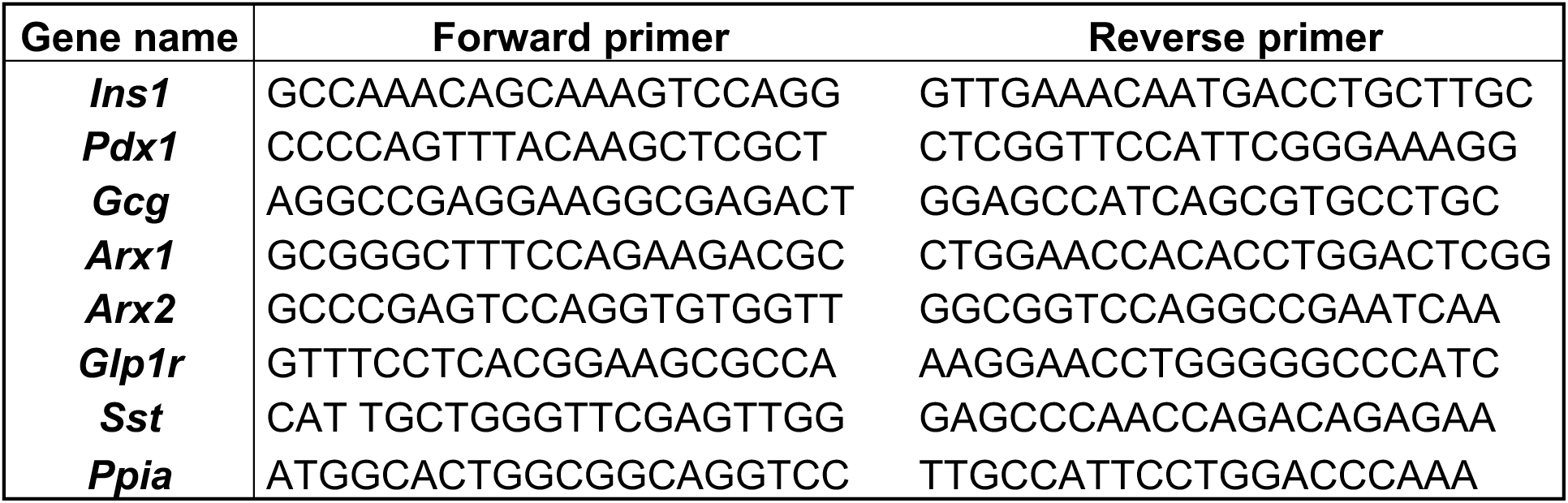
List of primers used in reverse transcription-quantitative PCR.

